# Adaptive divergence in the eyes of *Heliconius* butterflies likely contributes to pre- and post-mating isolation

**DOI:** 10.1101/2023.10.26.564160

**Authors:** Daniel Shane Wright, Juliana Rodriguez-Fuentes, Lisa Ammer, Kathy Darragh, Chi-Yun Kuo, W. Owen McMillan, Chris D. Jiggins, Stephen H. Montgomery, Richard M. Merrill

## Abstract

When populations experience different sensory conditions, natural selection may favor whole sensory system divergence, from the peripheral structures to the brain. We characterized the outer eye morphology of sympatric *Heliconius* species from different forest types, and their first-generation reciprocal hybrids to test for adaptive visual system divergence and hybrid disruption. In Panama, *Heliconius cydno* occurs in closed forests, whereas *Heliconius melpomene* resides in more open areas. Previous work has shown that, among wild individuals, *H. cydno* has larger eyes than *H. melpomene*, and there are heritable, habitat-associated differences in the visual brain structures that exceed neutral divergence expectations. Notably, hybrids have intermediate neural phenotypes, suggesting disruption. To test for similar effects in the visual periphery, we reared both species and their hybrids in common garden conditions. We confirm that *H. cydno* has larger eyes and provide new evidence that this is driven by selection. Hybrid eye morphology is more *H. melpomene*-like despite body size being intermediate, contrasting with neural trait intermediacy. Thus, eye morphology differences between *H. cydno* and *H. melpomene* are consistent with adaptive divergence, and when combined with previous neuroanatomy data, suggest hybrid visual system disruption due to mismatched patterns of intermediacy and dominance in the visual pathway.

## Introduction

Sensory systems mediate the transmission of information between an organism and its surroundings (Stevens, 2013). Natural selection is expected to favor divergent sensory phenotypes across populations exposed to different sensory conditions and/or which exploit different resources, potentially leading to both pre- and post-mating mating reproductive isolation (Dell’Aglio et al., 2023). Habitat-associated variation in sensory traits is well-documented, particularly for vision (Webster, 2015). However, most studies of visual adaptation between populations have focused on color vision in aquatic organisms (Carleton & Yourick, 2020; Cummings & Endler, 2018), whereas other aspects of visual perception are understudied. For example, we know little about how visual signals are evaluated (Rosenthal, 2018) or how adaptive processes operate at different levels within the visual pathway. Moreover, we lack information on the visual capabilities of hybrids despite their potential to contribute to reproductive isolation between species. Here, we characterize the outer eye morphology of *Heliconius* butterflies, examine evidence for adaptive divergence, and consider how this may lead to the disruption of visual systems in hybrids.

The compound eyes of insects represent an easily quantifiable sensory structure that directly affects visual perception. The insect compound eye consists of numerous independent photosensitive units, ommatidia, each of which receives visual (light) information and transfers it to the brain. Variation in the total number of ommatidia, their size, and density directly affects visual perception and often correlates with temporal activity (Greiner, 2006; Land, 1997; Stöckl et al., 2017; Warrant, 2004). For example, nocturnal and crepuscular species often have larger eyes and larger facets to enhance photon sensitivity in light-poor environments, as observed across insect taxa (Freelance et al., 2021). Associations between the local environment and the visual systems of diurnal insects are less studied and often included only as a comparison to other nocturnal species (e.g., Frederiksen & Warrant, 2008). Nonetheless, the conditions experienced by diurnal insects can vary greatly (Endler, 1993), and may represent important adaptations to the local environment.

*Heliconius* butterflies inhabit tropical and subtropical regions of the Americas and rely heavily on vision for foraging for both flowers and hostplants (Dell’Aglio et al., 2016; Gilbert, 1982), as well as finding and choosing suitable mates (Crane, 1955; Estrada & Jiggins, 2008; Hausmann et al., 2021; Jiggins et al., 2001; Merrill et al., 2019). In Panama, the closely related species *Heliconius melpomene* and *Heliconius cydno* are broadly sympatric, but occupy different forest types (Estrada & Jiggins, 2002). *H. melpomene* primarily lives in forest edge habitats, whereas *H. cydno* occurs deeper within the forests, with less light and increased habitat complexity (Fig. 1A) (DeVries, 1987; Estrada & Jiggins, 2002; Seymoure, 2016). Although patterns of opsin expression suggest few differences in wavelength sensitivity (McCulloch et al., 2017), recent data on brain morphologies of *H. melpomene* and *H. cydno* reported heritable differences in the size of the visual neuropils that exceed expected rates of neutral divergence (Montgomery et al., 2021). Using wild caught individuals, Seymoure et al. (2015) similarly found that i) *H. cydno* has larger eyes than *H. melpomene* and ii) that *H. cydno* males have larger eyes than *H. cydno* females (intraspecific differences in *H. melpomene* were non-significant). However, these results were based on individuals sampled as adults in their respective habitats and may include effects of environment-induced plasticity. Also, total ommatidia counts were measured for only two individuals for each species and sex, so statistical power to explore different eye morphology traits was limited.

**Figure 1.**
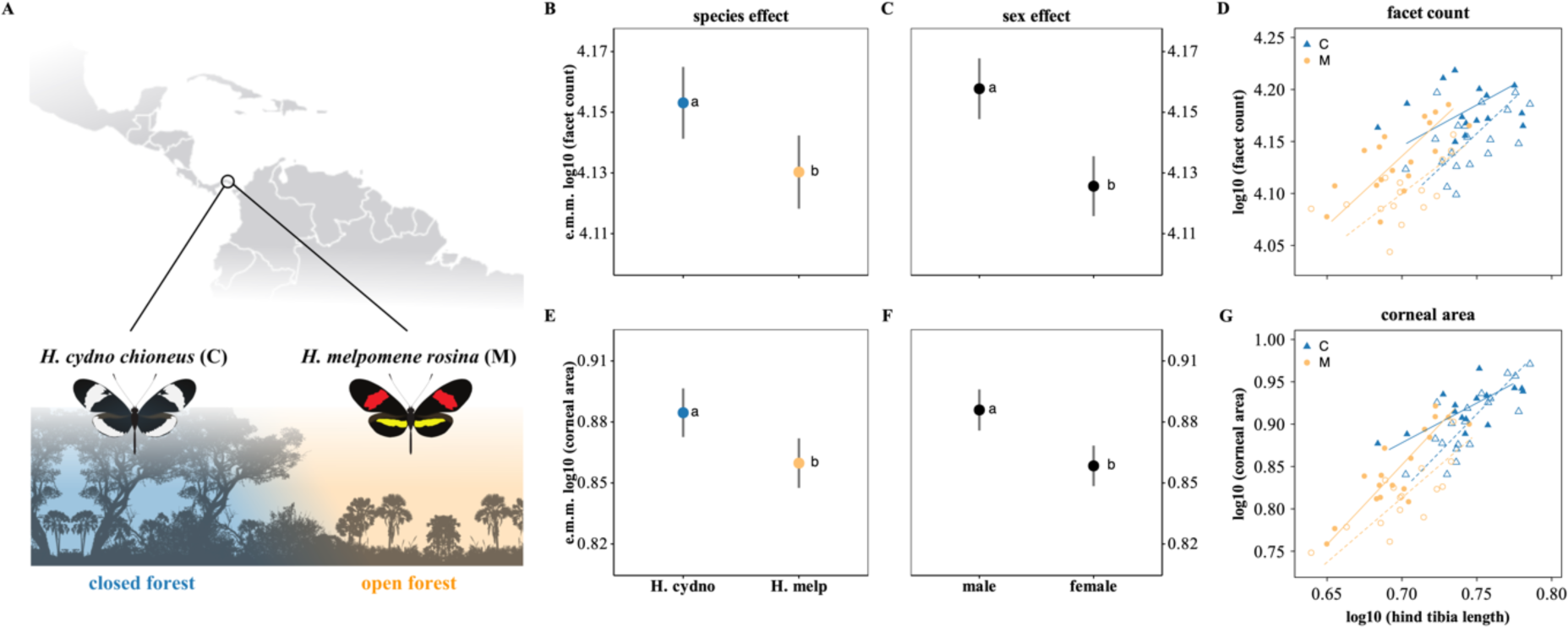
**(A)** *Heliconius cydno chioneus* and *Heliconius melpomene rosina* occur sympatrically in Panama but occupy different habitats: *H. cydno* is found in closed forest environments, whereas *H. melpomene* resides in open forests. **(B, C, E, F)** Estimated marginal means (e.m.m.) of the minimum adequate statistical models for **(B, C)** facet count and **(E, F)** corneal area, demonstrating the significant effects of **(B, E)** *species* and **(C, F)** *sex*, while accounting for other significant terms (*tibia length* and *sex/species*). The interaction between *species* and *sex* was never significant (FDR-p > 0.13). Different letters indicate significant differences (p < 0.05), and the error bars represent 95% confidence intervals. **(D, G)** Major axis regressions of **(D)** facet count or (**G)** corneal area and body size, measured as hind tibia length. Double-logarithmic plots are presented to explore the allometric relationships between eye morphology and body size. Males are represented by solid shapes and solid lines, and females are represented by open shapes and dashed lines. C = *H. cydno*; M = *H. melpomene*.

Given the evidence for selection acting on the visual processing regions of the brain, a more thorough examination of the visual periphery in these species is warranted. In particular, the role of plasticity and selection, and the potential for the mismatch of components of the visual system in interspecific hybrids have not yet been assessed. To address this, we characterized the outer eye morphology of *H. melpomene* and *H. cydno* to test for patterns of adaptive divergence in the visual system. First, we compared the eye morphology of butterflies (15+ for each species and sex) reared under common garden conditions. We then used a quantitative genetics approach to test if the species-specific eye differences are due to selection using *P_ST_*-*F_ST_* analysis. Finally, we report patterns of eye morphology in first-generation (F1) hybrids of *H. melpomene* and *H. cydno*.

## Methods

### Butterfly specimens

We established outbred stocks of *Heliconius cydno chioneus* (C) and *Heliconius melpomene rosina* (M) from butterflies caught in Gamboa and the nearby Soberanía National Park, Panama between 2007-2009 and 2015-2017. We generated reciprocal F1 hybrids between *H. cydno* and *H. melpomene* by either crossing a *H. cydno* female with *H. melpomene* a male (CxM), or a *H. melpomene* female with a *H. cydno* male (MxC). All pure individuals and hybrids were reared under common garden conditions in the Smithsonian Tropical Research Institute insectaries in Gamboa, and all specimens were preserved in DMSO/EDTA/NaCl and stored at −80° C as described in Merrill et al. (2019).

### Sample preparation

Samples were prepared following previously published methods (Seymoure et al., 2015; Wright et al., 2023). In brief, we thawed specimens at room temperature, and dissected out both eyes and the hind legs. The legs were immediately imaged (see below), while the eyes were placed in 20% sodium hydroxide (NaOH) for 18-24 hours to loosen the tissues behind the cuticular cornea. The following day, we cleaned each eye cuticle of excess tissue and mounted it on a microscope slide in Euparal (Carl Roth GmbH). The sample was left to dry overnight before imaging.

### Image analysis

We used ImageJ/Fiji (Schindelin et al., 2012) to analyze each mounted cornea for the total number of facets and total corneal area. All slides were imaged at 7.5x on a Leica M80 stereomicroscope fitted with a Leica Flexacam C1 camera and the *Leica Application Suite X* (LAS X) software. Each image contained a 1mm scale bar for calibration. Facet counts were measured via image thresholding and the *Analyze particles* function, and corneal surface area was measured with the *Freehand selection* and *Measure* options (full protocol provided as supplementary methods). This semi-automated method differs slightly from the approach used by Seymoure et al. (2015) but gives quantitatively similar results (Fig. S1). To account for differences in body size, we measured hind tibia length using the *Straight line* and *Measure* options. The number of facets (Pearson’s r [95% C.I.]: r (118) = 0.969 [0.956, 0.978]), corneal area (r (118) = 0.986 [0.979, 0.990]) and hind tibia length (r (101) = 0.930 [0.898, 0.952]) on the left vs. right sides of the butterfly were highly correlated. Therefore, for all subsequent analyses, we used only the left eye and left leg unless either was missing, damaged or had poor image quality (i.e., not all facets visible), then the right side was substituted.

### Statistical analysis

#### Eye morphology

We used linear models (lm function) in R to explore how facet count and corneal area are influenced by *species* (*H. cydno* vs. *H. melpomene*), *sex* (male vs. female), and *body size* (hind tibia length) as: *log10(facet count* or *corneal area) ∼ species * sex + log10(tibia length)*. Log10-transformations were used to normalize the residuals around the allometric relationships to meet the assumptions of normality (Thorpe, 1975). We also used linear models to assess i) the relationship between facet count and corneal as: *log10(corneal area) ∼ log10(facet count) * species * sex* and ii) body size differences (using hind tibia length as a body size proxy): *log10(tibia length) ∼ species + sex*. The significance of fixed effect parameters was determined by likelihood ratio tests via the *drop1* function, and minimum adequate models (MAM) were selected using statistical significance (Crawley, 2013; Nakagawa & Cuthill, 2007). We used the *Anova* function in the car package (Fox & Weisberg, 2018) to estimate significant fixed effect parameters and report false discovery rate (FDR; Benjamini & Hochberg, 1995) adjusted p-values (*p.adjust* function) to account for multiple testing. Model assumptions were confirmed via visual inspection (residual vs. fitted and normal Q-Q plots). To accurately visualize multiple significant fixed effects, we extracted and plotted the estimated marginal means from each MAM using the *emmeans* function in the emmeans package (Lenth et al., 2023).

We also explored whether the scaling relationships between eye morphology (facet count and corneal area) and body size (hind tibia length) differed for *H. cydno* and *H. melpomene* using major axis regressions via the *sma* function in the smatr package (Warton et al., 2012). Following the standard allometric scaling relationship, *log y = β log x + α*, we tested for shifts in the allometric slope (*β*). Where a common slope was supported, we subsequently tested for differences in *α* that would indicate ‘grade-shifts’ (test = “elevation”) and for major axis-shifts along the common slope (test = “shift”).

#### Test of selection

We next used a quantitative genetics approach to test whether eye morphology differences between *H. cydno* and *H. melpomene* are due to selection. *Q_ST_* is a quantitative genetic analogue of *F_ST_* that measures additive genetic variation among populations relative to total genetic variance. However, *Q_ST_* estimates for quantitative traits are difficult, so *Q_ST_* is often replaced with its phenotypic analogue *P_ST_* (Leinonen et al., 2013). Comparisons between *P_ST_* and *F_ST_* can be used as a test of divergent selection, where *P_ST_* values that exceed genome-wide *F_ST_* suggest greater phenotypic divergence than expected by neutral genetic divergence.

We calculated *P_ST_* values using the *Pst* function in the Pstat package (Silva & Silva, 2018) for raw, log10-transformed, and body-size corrected eye morphology measurements (i.e., facet count and corneal area). Allometrically scaled body-size correlations (using tibia length) per species and sex were performed via the *allomr* function in the allomr package (Schär, 2023). *P_ST_* approximation to *Q_ST_* depends on heritability, *h^2^*, and a scalar *c* that expresses the proportion of the total variance that is presumed to be due to additive genetic effects across populations (Brommer, 2011). Heritability estimates for facet count and corneal area are unknown for these species, so in addition to the default value of 1, we used varying *c/h^2^*ratios [ranging from 0.33 to 4, following Montgomery et al. (2021)]. Genome-wide *F_ST_* values between *H. c. chioneus* and *H. m. rosina* were obtained from Martin et al. (2013), derived from four wild-caught individuals per species using 100-kb genomic windows. We calculated p-values as the proportion of the *F_ST_* distribution that was above each *P_ST_* value (Leinonen et al., 2013); values above the 95^th^ percentile of the *F_ST_* distribution were interpreted as an indication of selection.

#### Hybrid phenotypes

We re-ran the linear models described above but included the CxM and MxC hybrids as two additional groups within the *species* factor. To test if hybrid body size was intermediate to *H. cydno* and *H. melpomene*, we also re-tested the linear model: *log10(tibia length) ∼ species + sex*. In the case of more than two categories per fixed effect parameter (i.e., *species*), we used post hoc Tukey tests (glht-multcomp package (Hothorn et al., 2008)) to obtain parameter estimates and report Bonferroni adjusted p-values for multiple comparisons.

## Results

### Heritable shifts in eye morphology between species residing in different forest types

In total, we sampled 15 male and 19 female *H. cydno* and 18 male and 15 female *H. melpomene*, all reared under common garden conditions. Using tibia length as a proxy for body size, we found that *H. cydno* was larger than *H. melpomene* (F_1, 65_ = 58.77, FDR-p < 0.001), but there was no evidence for sexual size dimorphism in either species (FDR-p = 0.7). All three fixed effects, *tibia length*, *species*, and *sex*, were retained in our model examining facet count. After accounting for size (larger butterflies had more facets: F_1, 63_ = 20.44, FDR-p < 0.001), *H. cydno* had more facets than *H. melpomene* (F_1, 63_ = 7.32, FDR-p = 0.013; Fig. 1B), and males had more facets than females (F_1, 63_ = 27.35, FDR-p < 0.001; Figs. 1C, S2). The interaction between *species* and *sex* was not significant (FDR-p = 0.7), implying conserved patterns of sexual dimorphism. We found similar results for corneal area (Fig. 1E, F), where all three fixed effects were also retained (Table S1). For all butterflies, larger corneal area was due to an increase in facet number, as evidenced by *facet count* significantly affecting *corneal area* in the model *log10(corneal area) ∼ log10(facet count) * species * sex* (F_1,64_ = 235.38, FDR-p < 0.001). Importantly, the *facet count x species* interaction was not significant in this model (FDR-p = 0.15), suggesting no differences in facet size between species.

Given the sex-specific differences in eye morphology reported above, we analyzed the scaling relationships between eye morphology and body size for males and females separately (Fig 1D, G). The only significant difference in slope (*β*) was when comparing the scaling relationship between corneal area and tibia length for *H. cydno* vs. *H. melpomene* males (FDR-p = 0.016; Table 1); all other comparisons were non-significant, confirming common slopes (FDR-p > 0.23; Table 1). In isolation, body size and facet count were uncorrelated for *H. cydno* males (r^2^ = 0.004, p = 0.8), but there was no statistical difference in scaling between the species suggesting this is potentially due to increased variance in *H. cydno* (Table 1). For all comparisons with a common slope, tests for grade-shifts (α) were non-significant (FDR-p > 0.4), but there was a significant shift along the common axis (FDR-p < 0.001; Table 1).

**Table 1.**
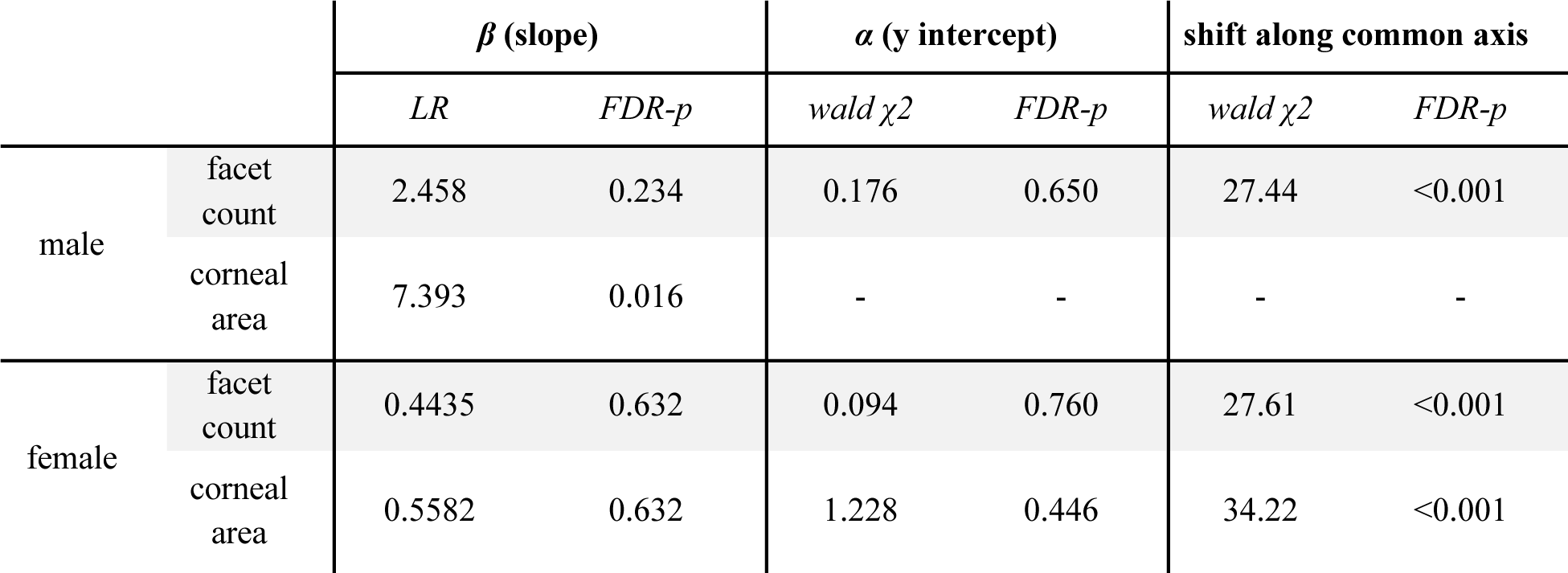
Scaling relationships between eye morphology and body size (hind tibia length) for *H. cydno* and *H. melpomene* males and females. No values are reported for male corneal area because tests for grade shifts (*α*) and shifts along the common axis are only appropriate with a common slope.

| | | $\beta$ (slope) | | $\alpha$ (y intercept) | | shift along common axis | |
| --- | --- | --- | --- | --- | --- | --- | --- |
|  |  | *LR* | *FDR-p* | *wald $\chi^2$* | *FDR-p* | *wald $\chi^2$* | *FDR-p* |
| male | facet count | 2.458 | 0.234 | 0.176 | 0.650 | 27.44 | <0.001 |
|  | corneal area | 7.393 | 0.016 | - | - | - | - |
| female | facet count | 0.4435 | 0.632 | 0.094 | 0.760 | 27.61 | <0.001 |
|  | corneal area | 0.5582 | 0.632 | 1.228 | 0.446 | 34.22 | <0.001 |

### Differences in eye morphology are driven by selection

Our *P_ST_*-*F_ST_* analyses suggest that the visual systems of *H. cydno* and *H. melpomene* have likely diverged as the result of selection rather than genetic drift. *P_ST_* was significantly higher than *F_ST_* for both facet count and corneal area (p < 0.001) for all comparisons (Fig. 2; Table S3), where the proportion of phenotypic variance due to additive genetic effects within-populations far exceeds the proportion of phenotypic variance due to additive genetic effects between-populations. Qualitatively similar results were obtained regardless of the phenotypic measurement evaluated (i.e., raw data, log10 transformed, allometrically corrected values; Table S3) and also when each sex was examined separately (Tables S4, S5).

**Figure 2.**
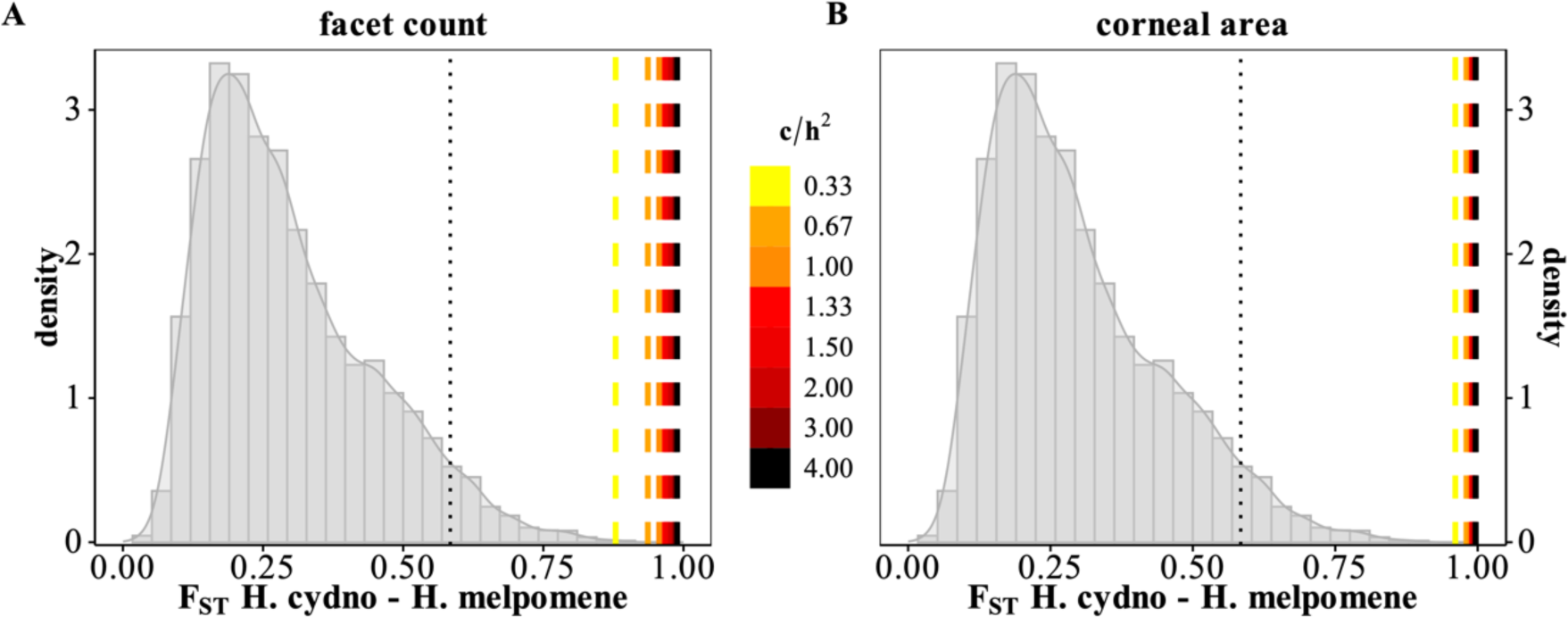
Location of the calculated *P_ST_* values for **(A)** facet count and **(B)** corneal area in the distribution of *F_ST_* values between *H. cydno chioneus* and *H. melpomene rosina* (values from Martin et al., 2013). Here, both morphological measurements are allometrically corrected using tibia length as a body-size proxy and presented using varying *c/h^2^* ratios (see Table S3 for *P_ST_* estimates using raw and log10 transformed values). The dotted line represents the 95^th^ percentile of the *F_ST_* distribution.

### Hybrid eye morphology is *H. melpomene*-like

We examined the eye morphology of 60 F1 hybrids, including 15 male and 15 female CxM individuals, and 15 female and 15 male MxC individuals (Table S2). Tukey post hoc (with Bonferroni correction) revealed patterns of intermediacy in body size (using tibia length as a body size proxy) for both hybrid types compared to the parental species: both CxM and MxC hybrids were larger than *H. melpomene* (p < 0.001), MxC was smaller than *H. cydno* (t = −3.148, p = 0.0124), and CxM did not differ from *H. cydno* (p = 1.0; Fig. 3A). However, F1 hybrid eye morphology was not intermediate to the parental species: both hybrid types had significantly fewer facets (p < 0.0035) than *H. cydno* but did not differ from *H. melpomene* (p = 1.0; Fig. 3B). Results for corneal area were similar (Table S2; Fig. S3, S4). These patterns were also evident when exploring the scaling relationships between eye morphology and body size of the F1 hybrids (Fig. 3C, D; Fig. S4).

**Figure 3.**
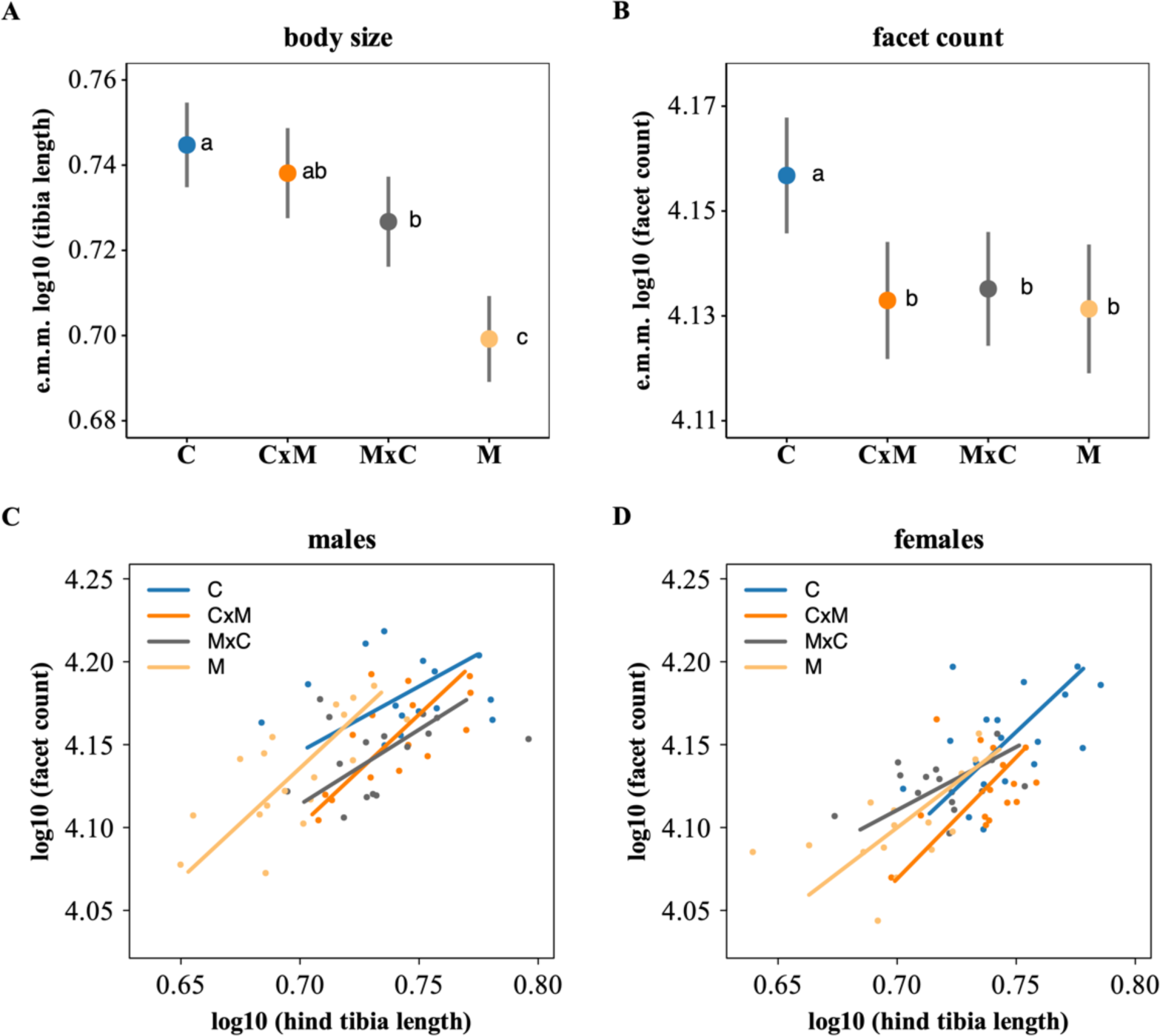
**(A)** Estimated marginal means (e.m.m.) of the minimum adequate statistical model for body size (using tibia length as a proxy) including F1 hybrids, demonstrating the significant effect of *species* (*sex* was non-significant, FDR-p = 0.48). E.m.m for **(B)** facet count, showing the significant effect of *species*, while accounting for significant *tibia length* and *sex* effects. In both plots, different letters indicate significant differences (Bonferroni adjusted p < 0.05), and the error bars represent 95% confidence intervals. **(C-D)** Major axis regressions of facet count and body size (tibia length) for males and females separately. Double-logarithmic plots are presented to explore the allometric relationships between eye morphology and body size. C = *H. cydno*; M = *H. melpomene*; CxM = F1 hybrid of *H. cydno* mother crossed with *H. melpomene* father; MxC = F1 hybrid of *H. melpomene* mother crossed with *H. cydno* father.

## Discussion

When populations are exposed to different sensory conditions, natural selection may favor divergence in sensory traits, which can contribute to speciation. We compared the outer eye morphology of *Heliconius* butterflies that occupy different environments to test for patterns of adaptive visual system divergence. Our results show that *H. cydno,* which occupies more visually complex closed-canopy forests, has larger eyes, and that this is a result of heritable differences in facet number. By combining our phenotypic data with genome wide estimates of *F_ST_*, we additionally provide strong evidence that selection has driven the divergence of eye morphology in these butterflies. Finally, we show that F1 hybrid eye morphology is not intermediate to the parental species, contrasting with patterns for body size and neural anatomy. This suggests that visual processing in hybrids may be disrupted by mismatches in different parts of the visual pathway.

Our results are consistent with previous work by Seymoure et al. (2015), which reported larger eyes for *H. cydno*, and bigger eyes in *H. cydno* (but not *H. melpomene*) males. However, the individuals sampled for our analyses were raised under common garden conditions, reducing the potential for environmental effects and genotype–environment interactions, which may give a distorted picture of the contribution of genetic variation on which selection can act (Brommer, 2011; Pujol et al., 2008). A potential caveat of our results is that we cut each cuticle four times to mount it on the microscope slides, which may have disrupted the semi-automated counts of individual facets. It is possible that more advanced 3D imaging techniques (e.g., Buffry et al., 2023), where cutting is not required, may give slightly higher total counts, though likely at the expense of overall sample size and associated statistical power. Regardless, our facet counts are consistent with prior work (Seymoure et al., 2015) (Fig. S1), and the close correspondence between wild and insectary-reared butterflies further suggests that the differences in eye morphology are largely heritable.

Habitat-associated variation in eye morphology has been reported across taxa (e.g., insects: (Greiner, 2006); mammals: (Veilleux & Lewis, 2011); fish: (Lisney et al., 2020); snakes: (Liu et al., 2012); primates: (Kirk, 2004)), and this variation is generally interpreted as an adaptive response to the local sensory conditions. For example, visual perception in insects is affected by the total number of ommatidia and their size/density (Greiner, 2006; Land, 1997; Warrant, 2004), and nocturnal and crepuscular species often possess larger eyes and larger facets to enhance photon sensitivity in low-light environments (Freelance et al., 2021). However, most studies do not formally evaluate the role of selection, which is a key element to define adaptation (Gould & Vrba, 1982). We are aware of only one study that has attempted to addresses this topic: Brandon et al. (2015) reported that eye size variation in a wild *Daphnia* population is associated with variation in fitness (reproductive output), suggesting that selection is operating, either directly or indirectly, on eye size variation (though the underlying mechanisms remain unknown).

In addition to revealing heritable differences in eye morphology, our results suggest that eye morphology in *H. cydno* and *H. melpomene* have evolved as the result of divergent selection, as opposed to genetic drift. Although our *P_ST_*-*F_ST_* approach to test for evidence of selection acting on eye morphology is limited by the difficulty in approximating *P_ST_* to *Q_ST_* (Brommer, 2011), these limitations are largely overcome by rearing our butterflies under common garden conditions (Leinonen et al., 2008, 2013). Moreover, as with most insects, the heritability of facet count and corneal area are unknown for the species used in this study. To account for this, we used a wide range of *c/h^2^* ratios, including the default assumption of *c = h^2^* (i.e., *c/h^2^* = 1), where the proportion of phenotypic variance due to additive genetic effects is the same for between-population variance and within-population variance (Brommer, 2011). In all cases, *P_ST_* values were higher than the 95^th^ percentile of the genome wide *F_ST_* distribution (Tables S3-S5).

In insects, eye size may increase due to an increase in the number of ommatidia, an increase in individual ommatidia size, or both. Our results suggest that larger eye size in *H. cydno*, and in males of both species, is predominantly due to an increase in ommatidia number. We did not measure facet diameter directly, but we found no effect of the *facet count x species* interaction on *corneal area*, suggesting no interspecific differences in facet size. Seymoure et al. (2015) measured facet diameter from a subset of ommatidia in six anatomical eye regions and reported larger facet diameter in wild *H. cydno* compared to *H. melpomene*, with no differences between sexes. However, these results stemmed from an analysis of covariance including other species (*H. sapho* and *H. erato*); there were no direct comparisons between *H. cydno* and *H. melpomene*. Future studies may benefit from exploring facet diameter more directly. Regardless, the larger eyes of *H. cydno,* and male *Heliconius*, appear to be largely due to an increase in ommatidia number. In insects, increased ommatidia number is thought to contribute to higher visual acuity (Land, 1997) - behavioral and morphological data revealing sexual dimorphism in the visual acuity of *H. erato* support this prediction (Wright et al., 2023). We note that our allometric analyses suggest part of the variation we observe is associated with body size, within and between species. However, our *P_ST_*-*F_ST_* analyses account for body size and still suggest a signal of selection. The pattern we observe in hybrids, where grade-shifts are clearly observed between *cydno* and *cydno x melpomene* hybrids (Figure 3C), also suggest a strong genetic component independent of body size. As such, while the behavioural impact of increased eye size likely depends on the raw numbers of facets, we conclude that selection on body size alone does not explain the increase in *cydno* eye size.

Multiple non-exclusive selective pressures could be driving the differences we report here. First, species differences may be explained by *H. cydno* occupying more complex closed-forest environments (Estrada & Jiggins, 2002), where more ommatidia are advantageous due to e.g., increased visual acuity (Land, 1997; Wright et al., 2023). More ommatidia in males may also be due to general ecological differences between the sexes, as males actively search for and identify mates (Rutowski, 2000; Yagi & Koyama, 1963). Interestingly, species differences persist for both sexes (Fig. S2) and when exploring the *P_ST_*-*F_ST_* results for each sex separately, we still observed evidence of selection (Tables S4, S5). Taken together, our results suggest that the selective pressure on males to have more ommatidia acts in both species in concert with selection for more ommatidia in closed-forest environments. Similar patterns may exist across *Heliconius*, but to date, few species have been surveyed for eye morphology.

Our results also mirror neuroanatomical comparisons between species across the *cydno-melpomene* clade, where larger visual neuropils are reported for *cydno*-clade species occupying closed-forest environments, as opposed to *H. melpomene* (Montgomery et al., 2021). These neural differences appear to be heritable adaptations, based on similar tests of selection to those reported here. Thus, the combined results on brain morphology (Montgomery et al., 2021) and those presented here suggest whole visual system adaptation, from the sensory periphery to the brain. Similar habitat-associated differences in neuroanatomy have been reported in other Neotropical butterflies (Montgomery & Merrill, 2017; Wainwright & Montgomery, 2022), indicating a broader pattern of sensory adaptation in ecologically divergent, but closely related, butterflies.

One notable difference in the results obtained for eye and neural traits, however, is in the pattern of variation among F1 interspecific hybrids. The eye morphology of *H. cydno* and *H. melpomene* F1 hybrids tended to be more *H. melpomene*-like. This contrasts with patterns for body size (Fig. 3A) and neuroanatomy, where hybrids are intermediate for at least some traits (Montgomery et al., 2021). The observation that hybrid eye morphology is *melpomene*-like, but hybrid body size tends to be intermediate indicates that these two traits have different underlying genetic architectures (i.e., eye size is not simply genetically correlated with increasing body size). Furthermore, evidence suggests that both eye morphology (this study) and neural anatomy (Montgomery et al., 2021) are under divergent selection and in the predicted direction (i.e., bigger eyes and larger visual neuropils in *H. cydno*). If these observations truly represent adaptations (as our results suggest), then hybrids may have suboptimal visual system functioning, whereby peripheral sensory structures (number of facets) are mismatched to subsequent processing regions (optic lobe neuropils). This would predict that hybrids suffer a fitness deficit due to lower performance in visually oriented tasks, such as foraging and mate detection, but behavioral experiments are required to test these predictions. In conclusion, the adaptive differences in eye structure we observe may contribute to ecologically based pre- and post-mating reproductive barriers.

## Acknowledgements

We thank Josephone Dessmann, Maria-Clara Melo, Moises Abanto, Adraia Tapia, Liz Evans for helping raise butterflies. We thank the Smithsonian Tropical Research Institute for providing research infrastructure, and the Autoridad Nacional del Ambiente for permission to collect butterflies in Panama (permit no. SE/A-22-09). This work was supported by a NERC Independent Research Fellowship (NE/N014936/1) to SHM, and a DFG Emmy Noether fellowship (Number: GZ: ME 4845/1-1) and an ERC starting grant (Grant number: 851040) to RMM.

## Data availability

The underlying data and R-scripts supporting the findings of this study are available at https://github.com/SpeciationBehaviour/Adaptive_divergence_Heliconius_eyes.

