## Supplementary material for "Adaptive divergence in the eyes of *Heliconius* butterflies likely contributes to pre- and post-mating isolation"

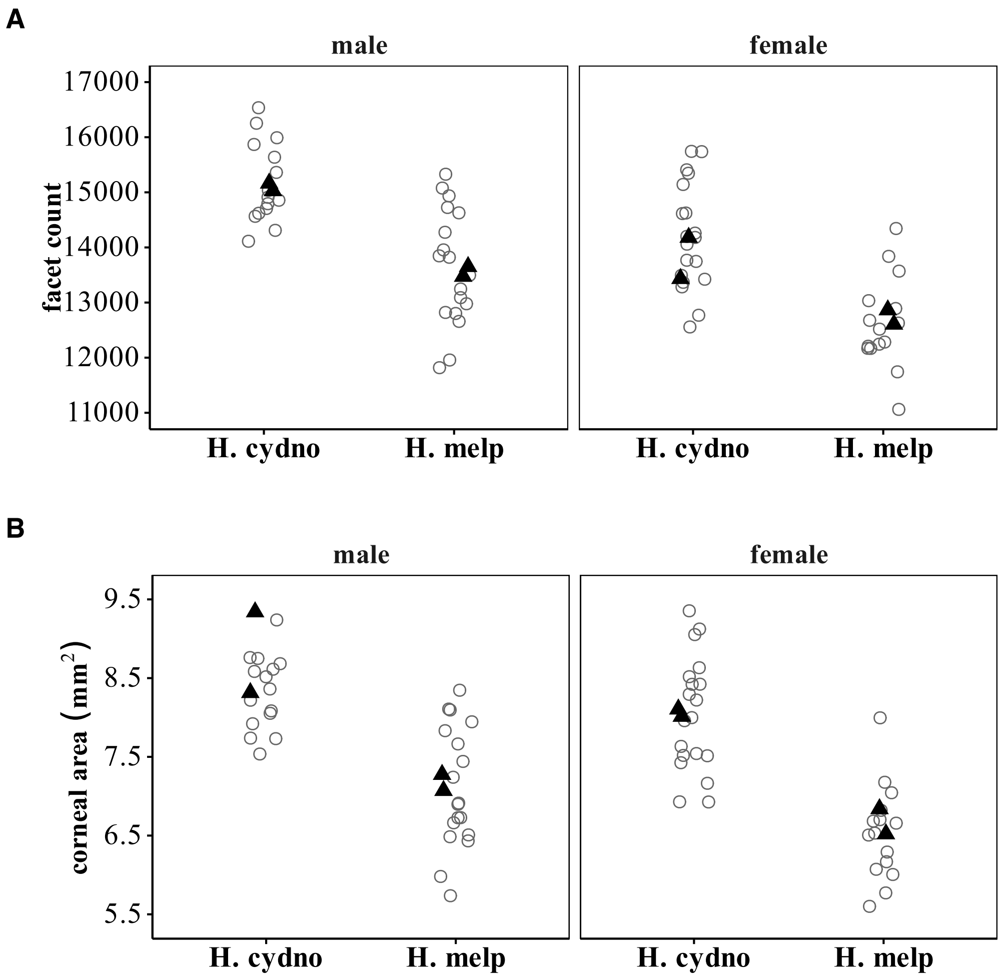


**Figure S1**. Total facet count and corneal area from this study (open circles) and Seymoure et al. (2015; black triangles) are quantitatively similar despite slightly different image analysis methods. Values from Figure 7 in Seymoure et al. (2015) were extracted using *WebPlotDigitizer* (https://automeris.io/WebPlotDigitizer/).

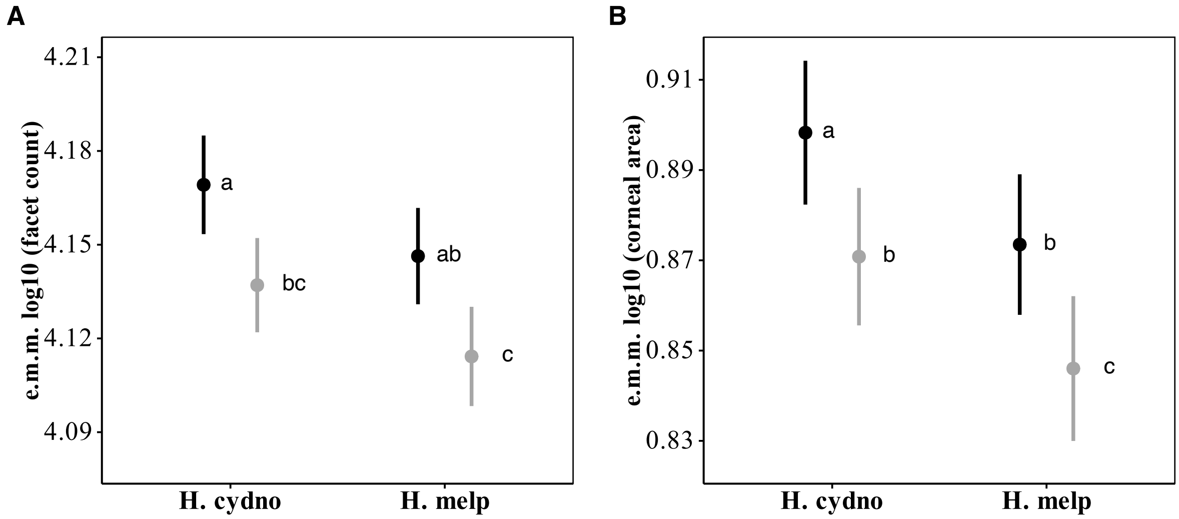


**Figure S2**. Estimated marginal means (e.m.m.) of the minimum adequate statistical models for **(A)** facet count and **(B)** corneal area, demonstrating the non-significant (FDR-p > 0.14) interaction between *species* and *sex*, while accounting for the significant effect of *tibia length*. Different letters indicate pairwise significant differences (Bonferroni adjusted $p < 0.05$), and the error bars represent 95% confidence intervals. Males are in black, and females are in grey.

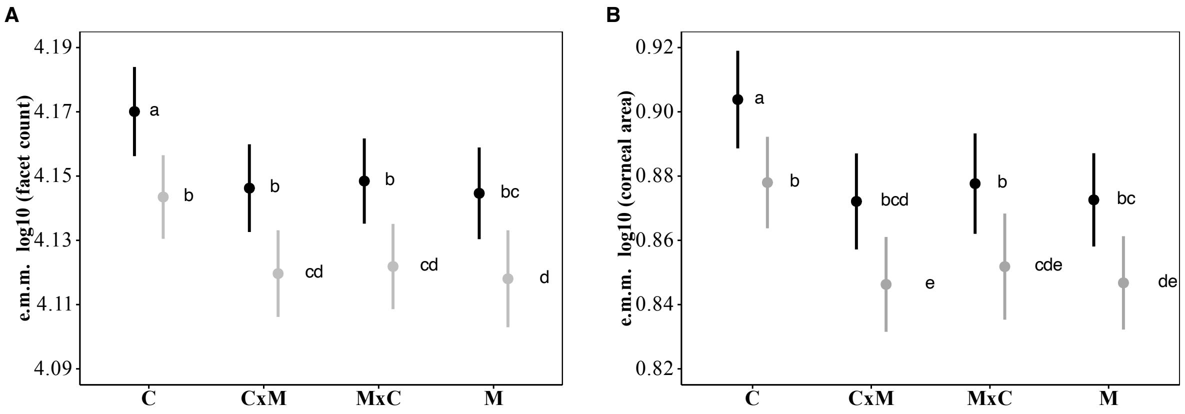


**Figure S3**. Estimated marginal means (e.m.m.) of the minimum adequate statistical models for **(A)** facet count and **(B)** corneal area including reciprocal F1 hybrids, demonstrating the non-significant (FDR-p > 0.29) interaction between *species* and *sex*, while accounting for the significant effect of *tibia length*. Different letters indicate pairwise significant differences (Bonferroni adjusted p < 0.05), and the error bars represent 95% confidence intervals. Males are in black, and females are in grey. C = *H. cydno*; M = *H. melpomene*; CxM = F1 hybrid of *H. cydno* mother crossed with *H. melpomene* father; MxC = F1 hybrid of *H. melpomene* mother crossed with *H. cydno* father.

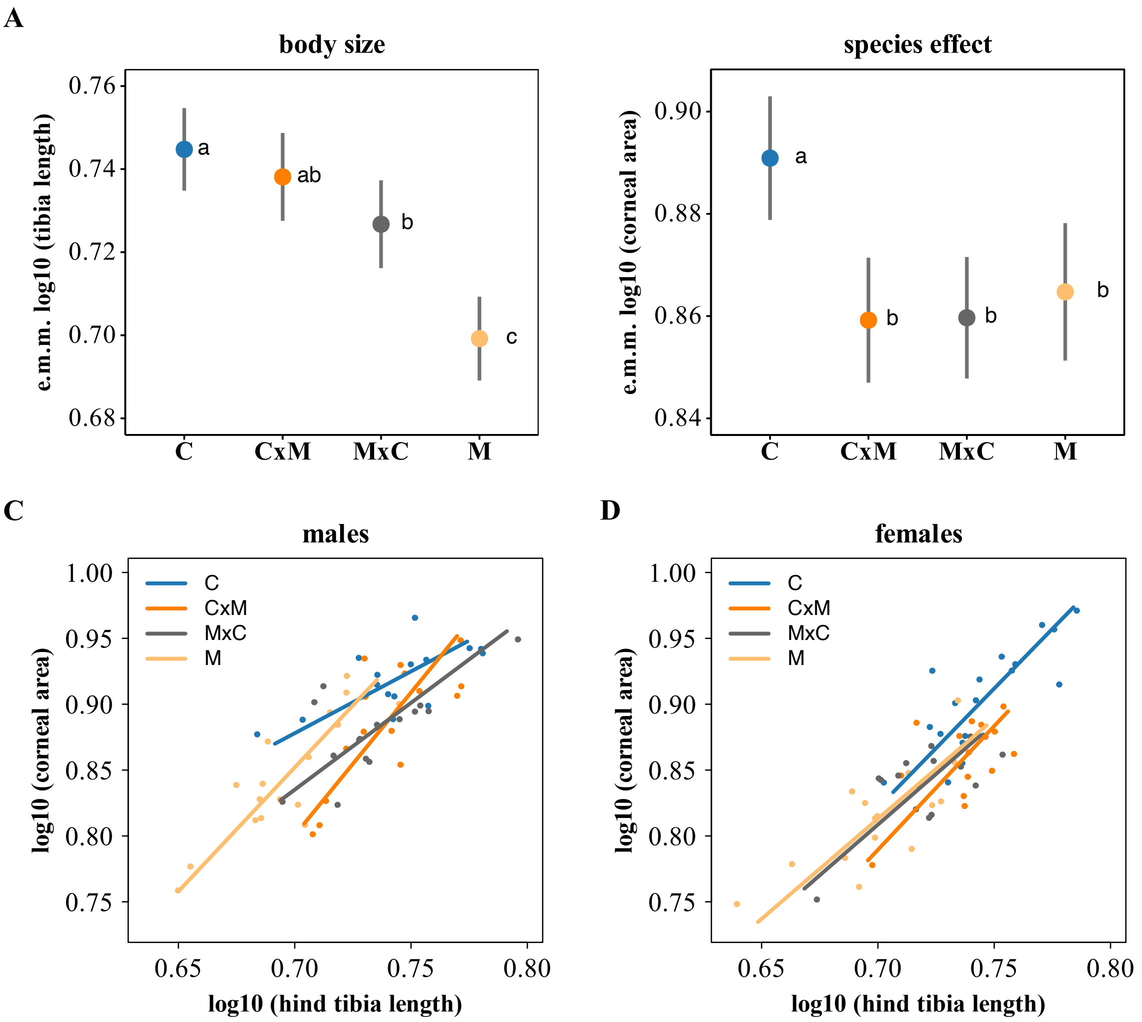


**Figure S4**. **(A)** Estimated marginal means (e.m.m.) of the minimum adequate statistical model for body size (using hind tibia length as a proxy) including F1 hybrids, demonstrating the significant effect of *species* (*sex* was non-significant, FDR-p = 0.44). E.m.m for **(B)** corneal area, showing the significant effect of *species*, while accounting for significant *tibia length* and *sex* effects. In both plots, different letters indicate significant differences (Bonferroni adjusted $p < 0.05$), and the error bars represent 95% confidence intervals. **(C-D)** Major axis regressions of corneal area and body size (hind tibia length) for males and females separately. Double-logarithmic plots are presented to explore the allometric relationships between eye morphology and body size. C = *H. cydno*; M = *H. melpomene*; CxM = F1 hybrid of *H. cydno* mother crossed with *H. melpomene* father; MxC = F1 hybrid of *H. melpomene* mother crossed with *H. cydno* father.

| | facet count | | | corneal area | | |
|---|---|---|---|---|---|---|
| **fixed effects** | **F value** | **df (resid df)** | **FDR-p** | **F value** | **df (resid df)** | **FDR-p** |
| **species** | 7.317 | 1 (63) | 0.013 | 8.469 | 1 (63) | 0.008 |
| **sex** | 27.346 | 1 (63) | <0.001 | 19.618 | 1 (63) | <0.001 |
| **tibia length** | 20.444 | 1 (63) | <0.001 | 90.417 | 1 (63) | <0.001 |
| **species * sex** | 0.131 | 1 (62) | 0.753 | 2.749 | 1 (62) | 0.141 |

**Table S1**. Fixed effect parameter estimates from linear models exploring facet count and corneal area for *H. cydno* and *H. melpomene*. For both models, log10 transformations of facet count/corneal area and tibia length were used to normalize the residuals around the allometric relationships to meet the assumptions of normality. False discovery rate adjusted p-values are reported to account for multiple hypothesis testing.

| | facet count | | | corneal area | | |
|---|---|---|---|---|---|---|
| **fixed effects** | **F value** | **df (resid df)** | **FDR-p** | **F value** | **df (resid df)** | **FDR-p** |
| **species** | 7.1147 | 3 (121) | <0.001 | 10.333 | 3 (121) | <0.001 |
| **sex** | 40.4010 | 1 (121) | <0.001 | 31.791 | 1 (121) | <0.001 |
| **tibia length** | 29.7570 | 1 (121) | <0.001 | 137.413 | 1 (121) | <0.001 |
| **species * sex** | 1.4072 | 3 (118) | 0.298 | 0.9002 | 3 (118) | 0.488 |

**Table S2**. Fixed effect parameter estimates from linear models exploring facet count and corneal area for *H. cydno* (C), *H. melpomene* (M), and their reciprocal F1 hybrids (CxM and MxC). For both models, log10 transformations of facet count/corneal area and tibia length were used to normalize the residuals around the allometric relationships to meet the assumptions of normality. False discovery rate adjusted p-values are reported to account for multiple hypothesis testing.

| morphological measure | measurement type | $c/h^2$ | $P_{ST}$ | Lower bound 95% CI | Upper bound 95% CI | p-value |
|---|---|---|---|---|---|---|
| facet count | raw value | 0.33 | 0.8457 | 0.6906 | 0.9250 | 0.0006 |
| | | 0.67 | 0.9176 | 0.8320 | 0.9578 | 0.0000 |
| | | 1.00 | 0.9432 | 0.8709 | 0.9718 | 0.0000 |
| | | 1.33 | 0.9567 | 0.9002 | 0.9790 | 0.0000 |
| | | 1.50 | 0.9614 | 0.9150 | 0.9810 | 0.0000 |
| | | 2.00 | 0.9708 | 0.9339 | 0.9858 | 0.0000 |
| | | 3.00 | 0.9803 | 0.9544 | 0.9901 | 0.0000 |
| | | 4.00 | 0.9852 | 0.9652 | 0.9929 | 0.0000 |
| | log10 transformed | 0.33 | 0.8463 | 0.6987 | 0.9218 | 0.0006 |
| | | 0.67 | 0.9179 | 0.8344 | 0.9569 | 0.0000 |
| | | 1.00 | 0.9435 | 0.8784 | 0.9734 | 0.0000 |
| | | 1.33 | 0.9569 | 0.9055 | 0.9788 | 0.0000 |
| | | 1.50 | 0.9616 | 0.9066 | 0.9814 | 0.0000 |
| | | 2.00 | 0.9709 | 0.9303 | 0.9858 | 0.0000 |
| | | 3.00 | 0.9804 | 0.9542 | 0.9908 | 0.0000 |
| | | 4.00 | 0.9852 | 0.9622 | 0.9927 | 0.0000 |
| | allometrically corrected | 0.33 | 0.8789 | 0.7812 | 0.9295 | 0.0004 |
| | | 0.67 | 0.9364 | 0.8820 | 0.9652 | 0.0000 |
| | | 1.00 | 0.9565 | 0.9168 | 0.9752 | 0.0000 |
| | | 1.33 | 0.9669 | 0.9327 | 0.9820 | 0.0000 |
| | | 1.50 | 0.9706 | 0.9433 | 0.9837 | 0.0000 |
| | | 2.00 | 0.9778 | 0.9530 | 0.9881 | 0.0000 |
| | | 3.00 | 0.9851 | 0.9724 | 0.9919 | 0.0000 |
| | | 4.00 | 0.9888 | 0.9786 | 0.9937 | 0.0000 |
| corneal area | raw value | 0.33 | 0.9095 | 0.8391 | 0.9496 | 0.0000 |
| | | 0.67 | 0.9533 | 0.9139 | 0.9745 | 0.0000 |
| | | 1.00 | 0.9682 | 0.9406 | 0.9829 | 0.0000 |
| | | 1.33 | 0.9759 | 0.9564 | 0.9873 | 0.0000 |
| | | 1.50 | 0.9786 | 0.9611 | 0.9887 | 0.0000 |
| | | 2.00 | 0.9838 | 0.9706 | 0.9911 | 0.0000 |
| | | 3.00 | 0.9892 | 0.9796 | 0.9944 | 0.0000 |
| | | 4.00 | 0.9919 | 0.9845 | 0.9958 | 0.0000 |
| | log10 transformed | 0.33 | 0.9091 | 0.8416 | 0.9504 | 0.0000 |
| | | 0.67 | 0.9531 | 0.9167 | 0.9742 | 0.0000 |
| | | 1.00 | 0.9681 | 0.9386 | 0.9836 | 0.0000 |
| | | 1.33 | 0.9758 | 0.9540 | 0.9866 | 0.0000 |
| | | 1.50 | 0.9785 | 0.9618 | 0.9885 | 0.0000 |
| | | 2.00 | 0.9838 | 0.9698 | 0.9914 | 0.0000 |
| | | 3.00 | 0.9891 | 0.9788 | 0.9942 | 0.0000 |
| | | 4.00 | 0.9918 | 0.9849 | 0.9957 | 0.0000 |
| | allometrically corrected | 0.33 | 0.9608 | 0.9433 | 0.9758 | 0.0000 |
| | | 0.67 | 0.9803 | 0.9707 | 0.9876 | 0.0000 |
| | | 1.00 | 0.9867 | 0.9797 | 0.9919 | 0.0000 |
| | | 1.33 | 0.9900 | 0.9846 | 0.9940 | 0.0000 |
| | | 1.50 | 0.9911 | 0.9862 | 0.9946 | 0.0000 |
| | | 2.00 | 0.9933 | 0.9892 | 0.9959 | 0.0000 |
| | | 3.00 | 0.9955 | 0.9932 | 0.9973 | 0.0000 |
| | | 4.00 | 0.9966 | 0.9948 | 0.9979 | 0.0000 |

**Table S3**. Calculated $P_{ST}$ values and the corresponding confidence intervals for facet count and corneal area using different $c/h^2$ ratios. P-values were calculated as the proportion of the $F_{ST}$ distribution above each $P_{ST}$ value. Allometric corrections use tibia length as a body size proxy.

| morphological measure | measurement type | $c/h^2$ | $P_{ST}$ | Lower bound 95% CI | Upper bound 95% CI | p-value |
|---|---|---|---|---|---|---|
| facet count | raw value | 0.33 | 0.7895 | 0.5866 | 0.9009 | 0.0033 |
| | | 0.67 | 0.8839 | 0.7515 | 0.9475 | 0.0000 |
| | | 1.00 | 0.9191 | 0.8149 | 0.9659 | 0.0000 |
| | | 1.33 | 0.9379 | 0.8552 | 0.9733 | 0.0000 |
| | | 1.50 | 0.9446 | 0.8672 | 0.9761 | 0.0000 |
| | | 2.00 | 0.9578 | 0.8931 | 0.9809 | 0.0000 |
| | | 3.00 | 0.9715 | 0.9269 | 0.9877 | 0.0000 |
| | | 4.00 | 0.9785 | 0.9452 | 0.9908 | 0.0000 |
| | log10 transformed | 0.33 | 0.7856 | 0.5706 | 0.8970 | 0.0038 |
| | | 0.67 | 0.8815 | 0.7426 | 0.9455 | 0.0002 |
| | | 1.00 | 0.9174 | 0.8122 | 0.9643 | 0.0000 |
| | | 1.33 | 0.9366 | 0.8588 | 0.9732 | 0.0000 |
| | | 1.50 | 0.9433 | 0.8679 | 0.9751 | 0.0000 |
| | | 2.00 | 0.9569 | 0.8955 | 0.9821 | 0.0000 |
| | | 3.00 | 0.9708 | 0.9283 | 0.9872 | 0.0000 |
| | | 4.00 | 0.9780 | 0.9472 | 0.9909 | 0.0000 |
| | allometrically corrected | 0.33 | 0.8564 | 0.7568 | 0.9190 | 0.0006 |
| | | 0.67 | 0.9237 | 0.8619 | 0.9600 | 0.0000 |
| | | 1.00 | 0.9476 | 0.9069 | 0.9706 | 0.0000 |
| | | 1.33 | 0.9601 | 0.9283 | 0.9786 | 0.0000 |
| | | 1.50 | 0.9644 | 0.9351 | 0.9822 | 0.0000 |
| | | 2.00 | 0.9731 | 0.9510 | 0.9853 | 0.0000 |
| | | 3.00 | 0.9819 | 0.9665 | 0.9905 | 0.0000 |
| | | 4.00 | 0.9864 | 0.9758 | 0.9931 | 0.0000 |
| corneal area | raw value | 0.33 | 0.8253 | 0.6739 | 0.9205 | 0.0016 |
| | | 0.67 | 0.9056 | 0.7997 | 0.9598 | 0.0000 |
| | | 1.00 | 0.9347 | 0.8684 | 0.9737 | 0.0000 |
| | | 1.33 | 0.9501 | 0.8872 | 0.9790 | 0.0000 |
| | | 1.50 | 0.9555 | 0.8987 | 0.9811 | 0.0000 |
| | | 2.00 | 0.9663 | 0.9235 | 0.9854 | 0.0000 |
| | | 3.00 | 0.9772 | 0.9493 | 0.9905 | 0.0000 |
| | | 4.00 | 0.9828 | 0.9602 | 0.9926 | 0.0000 |
| | log10 transformed | 0.33 | 0.8174 | 0.6597 | 0.9193 | 0.0017 |
| | | 0.67 | 0.9009 | 0.8161 | 0.9597 | 0.0000 |
| | | 1.00 | 0.9314 | 0.8595 | 0.9712 | 0.0000 |
| | | 1.33 | 0.9475 | 0.8792 | 0.9779 | 0.0000 |
| | | 1.50 | 0.9532 | 0.8979 | 0.9802 | 0.0000 |
| | | 2.00 | 0.9645 | 0.9174 | 0.9843 | 0.0000 |
| | | 3.00 | 0.9760 | 0.9477 | 0.9898 | 0.0000 |
| | | 4.00 | 0.9819 | 0.9604 | 0.9926 | 0.0000 |
| | allometrically corrected | 0.33 | 0.9398 | 0.9037 | 0.9676 | 0.0000 |
| | | 0.67 | 0.9694 | 0.9501 | 0.9846 | 0.0000 |
| | | 1.00 | 0.9793 | 0.9663 | 0.9900 | 0.0000 |
| | | 1.33 | 0.9843 | 0.9732 | 0.9920 | 0.0000 |
| | | 1.50 | 0.9861 | 0.9766 | 0.9931 | 0.0000 |
| | | 2.00 | 0.9895 | 0.9826 | 0.9948 | 0.0000 |
| | | 3.00 | 0.9930 | 0.9882 | 0.9966 | 0.0000 |
| | | 4.00 | 0.9947 | 0.9911 | 0.9974 | 0.0000 |

**Table S4**. Calculated $P_{ST}$ values and the corresponding confidence intervals for male facet count and corneal area using different $c/h^2$ ratios. P-values were calculated as the proportion of the $F_{ST}$ distribution above each $P_{ST}$ value. Allometric corrections use tibia length as a body size proxy.

| morphological measure | measurement type | $c/h^2$ | $P_{ST}$ | Lower bound 95% CI | Upper bound 95% CI | p-value |
|---|---|---|---|---|---|---|
| facet count | raw value | 0.33 | 0.8065 | 0.6100 | 0.9063 | 0.0019 |
| facet count | raw value | 0.67 | 0.8943 | 0.7441 | 0.9533 | 0.0000 |
| facet count | raw value | 1.00 | 0.9266 | 0.8037 | 0.9696 | 0.0000 |
| facet count | raw value | 1.33 | 0.9438 | 0.8587 | 0.9771 | 0.0000 |
| facet count | raw value | 1.50 | 0.9499 | 0.8662 | 0.9770 | 0.0000 |
| facet count | raw value | 2.00 | 0.9619 | 0.9081 | 0.9833 | 0.0000 |
| facet count | raw value | 3.00 | 0.9743 | 0.9288 | 0.9892 | 0.0000 |
| facet count | raw value | 4.00 | 0.9806 | 0.9495 | 0.9921 | 0.0000 |
| facet count | log10 transformed | 0.33 | 0.8108 | 0.6138 | 0.9131 | 0.0018 |
| facet count | log10 transformed | 0.67 | 0.8969 | 0.7592 | 0.9527 | 0.0000 |
| facet count | log10 transformed | 1.00 | 0.9285 | 0.8327 | 0.9699 | 0.0000 |
| facet count | log10 transformed | 1.33 | 0.9453 | 0.8750 | 0.9771 | 0.0000 |
| facet count | log10 transformed | 1.50 | 0.9512 | 0.8795 | 0.9784 | 0.0000 |
| facet count | log10 transformed | 2.00 | 0.9629 | 0.8999 | 0.9846 | 0.0000 |
| facet count | log10 transformed | 3.00 | 0.9750 | 0.9239 | 0.9889 | 0.0000 |
| facet count | log10 transformed | 4.00 | 0.9811 | 0.9499 | 0.9924 | 0.0000 |
| facet count | allometrically corrected | 0.33 | 0.8500 | 0.7249 | 0.9219 | 0.0006 |
| facet count | allometrically corrected | 0.67 | 0.9200 | 0.8508 | 0.9618 | 0.0000 |
| facet count | allometrically corrected | 1.00 | 0.9450 | 0.8970 | 0.9717 | 0.0000 |
| facet count | allometrically corrected | 1.33 | 0.9581 | 0.9182 | 0.9780 | 0.0000 |
| facet count | allometrically corrected | 1.50 | 0.9626 | 0.9295 | 0.9813 | 0.0000 |
| facet count | allometrically corrected | 2.00 | 0.9717 | 0.9446 | 0.9866 | 0.0000 |
| facet count | allometrically corrected | 3.00 | 0.9810 | 0.9645 | 0.9908 | 0.0000 |
| facet count | allometrically corrected | 4.00 | 0.9857 | 0.9713 | 0.9928 | 0.0000 |
| corneal area | raw value | 0.33 | 0.8721 | 0.7545 | 0.9385 | 0.0005 |
| corneal area | raw value | 0.67 | 0.9326 | 0.8643 | 0.9681 | 0.0000 |
| corneal area | raw value | 1.00 | 0.9538 | 0.9043 | 0.9781 | 0.0000 |
| corneal area | raw value | 1.33 | 0.9649 | 0.9289 | 0.9837 | 0.0000 |
| corneal area | raw value | 1.50 | 0.9687 | 0.9336 | 0.9863 | 0.0000 |
| corneal area | raw value | 2.00 | 0.9764 | 0.9500 | 0.9888 | 0.0000 |
| corneal area | raw value | 3.00 | 0.9841 | 0.9659 | 0.9927 | 0.0000 |
| corneal area | raw value | 4.00 | 0.9880 | 0.9738 | 0.9945 | 0.0000 |
| corneal area | log10 transformed | 0.33 | 0.8783 | 0.7730 | 0.9402 | 0.0005 |
| corneal area | log10 transformed | 0.67 | 0.9361 | 0.8759 | 0.9696 | 0.0000 |
| corneal area | log10 transformed | 1.00 | 0.9563 | 0.9094 | 0.9796 | 0.0000 |
| corneal area | log10 transformed | 1.33 | 0.9668 | 0.9285 | 0.9843 | 0.0000 |
| corneal area | log10 transformed | 1.50 | 0.9704 | 0.9402 | 0.9864 | 0.0000 |
| corneal area | log10 transformed | 2.00 | 0.9777 | 0.9520 | 0.9898 | 0.0000 |
| corneal area | log10 transformed | 3.00 | 0.9850 | 0.9675 | 0.9931 | 0.0000 |
| corneal area | log10 transformed | 4.00 | 0.9887 | 0.9740 | 0.9946 | 0.0000 |
| corneal area | allometrically corrected | 0.33 | 0.9472 | 0.9089 | 0.9729 | 0.0000 |
| corneal area | allometrically corrected | 0.67 | 0.9733 | 0.9547 | 0.9875 | 0.0000 |
| corneal area | allometrically corrected | 1.00 | 0.9819 | 0.9690 | 0.9915 | 0.0000 |
| corneal area | allometrically corrected | 1.33 | 0.9864 | 0.9760 | 0.9931 | 0.0000 |
| corneal area | allometrically corrected | 1.50 | 0.9879 | 0.9791 | 0.9942 | 0.0000 |
| corneal area | allometrically corrected | 2.00 | 0.9909 | 0.9843 | 0.9957 | 0.0000 |
| corneal area | allometrically corrected | 3.00 | 0.9939 | 0.9894 | 0.9973 | 0.0000 |
| corneal area | allometrically corrected | 4.00 | 0.9954 | 0.9919 | 0.9978 | 0.0000 |

**Table S5**. Calculated $P_{ST}$ values and the corresponding confidence intervals for female facet count and corneal area using different $c/h^2$ ratios. P-values were calculated as the proportion of the $F_{ST}$ distribution above each $P_{ST}$ value. Allometric corrections use tibia length as a body size proxy.
